# Helix-to-Beta-Sheet Transition Drives Self-Assembly of Glutamate Transporter EAA1 Splice Peptides

**DOI:** 10.1101/2025.08.13.670123

**Authors:** Alper Karagöl, Taner Karagöl

## Abstract

Truncated isoforms play a critical role in understanding the structural and functional properties of membrane proteins, including glutamate transporters. Here, we molecularly characterize two helical truncated isoforms of the human glutamate transporter EAA1. Using an integrative multi-omics and computational approach, we show that these isoforms, particularly one derived from the N-terminus, do not adopt the canonical transporter fold. Instead, they self-assemble into stable, β-sheet-enriched oligomers, a structure previously unobserved for this protein family. Furthermore, we identified a water-soluble truncated isoform (A0A7P0TAF5) of the membranous canonical EAA1, revealing that self-assembly is not confined to membranous isoforms of EAA1. This finding uncovers a previously unrecognized functional class of truncated isoforms capable of initiating assembly in the soluble state. Our 500ns molecular dynamics simulations further reveal that the truncation alters the native conformational dynamics, promoting a transition into semi-helical β-structures over time. In a model bilayer, β-Sheet-driven octamerization of the helical EAA1 isoform A0A7P0Z4F7 induces localized upper leaflet membrane pitting during 250ns all-atom simulation. Helix to β-sheet oligomer transitions is a known pathological hallmark of neurodegenerative disorders such as Alzheimer’s disease. Our findings thus uncover a potential new mechanism for glutamate transporter involvement in neurodegeneration and identify the N-terminal domain as a promising therapeutic target. This work highlights how alternative splicing can generate isoforms with novel interaction patterns and distinct molecular conformations.

## Introduction

The synaptic effects of glutamate are terminated by the action of excitatory amino acid transporters (EAATs) located on the plasma membrane of astrocytes and neurons [1,2,3]. This reuptake helps to prevent the excitotoxicity [1,2,3]. Thus, dysregulations of glutamate transporter’s function have been linked to the etiologies of many conditions, including but not limited to, neurodegenerative disease [2], affective disorders [4]and schizophrenia [3,4,5].

The molecular complexity of these transporters extends beyond their canonical full-length structures, with alternative splicing and protein truncation generating diverse isoforms that challenge our understanding of protein structure and function [6,7,8]. Alternative splicing, initiation and promoter usage introduce a re-distribution in the coding regions of mRNA transcripts (exons) [6], leads to the generation of multiple isoforms for a single transporter gene, in this case *SLC1A3*. While previous research has primarily focused on complete protein configurations [7,8], emerging evidence suggests that truncated isoforms may possess unique structural and functional characteristics that remain largely unexplored [7,8]. For instance, expression of EAA1 and EAA2 splice variants is significantly altered in enriched populations of ACC pyramidal neurons in postmortem brains of individuals with schizophrenia [5]. Truncated isoforms lack specific exons for transport process but may produce some modulatory effect by heteromerization [8].

Our previous studies uncovered their potential effects as inhibitors of canonical assemblies, according to molecular dynamics simulations followed by complex structure predictions [8]. While their inhibitor potential was discovered, some isoforms further showed a tendency to form stable dimers in membranous systems [8]. Notably, all self-assembling EAAT isoforms identified to date were insoluble in water [8]. However, the precise functions of these assemblies, along with the dynamic behaviour of soluble truncated isoforms, remain hidden. Truncation events are known to induce significant alterations in protein conformation, potentially giving rise to novel molecular assemblies with distinct biophysical attributes [9,10]. An example is the Alzheimer β-peptide, which shifts between left-handed 3□-helix, β-strand, and random coil structures depending on the temperature [10]. Sheet-helical transformations describe the remarkable ability of certain specifically designed peptides to switch between stable β-sheet and α-helical secondary structures [11,12]. These peptides, often termed ionic self-complementary oligopeptides, typically possess alternating hydrophobic and hydrophilic residues, allowing for stable β-sheet formation through hydrophobic interactions on one face and extensive ionic interactions on the other [11]. Zhang and Rich previously discovered that a 16-amino acid oligopeptide adopts a stable β-sheet structure in water and self-assembles into a macroscopic matrix in physiological conditions, the peptide undergoes a sharp conformational shift from a β-sheet to a stable α-helix, bypassing any detectable random-coil intermediate [12]. These simple, versatile systems mark a major step in molecular engineering for diverse applications [12,13].

Understanding these alternative protein configurations is paramount for comprehending the mechanisms underlying neural signaling and potential dysregulation in neurological disorders [11,12,13]. To this end, multi-omics analysis combined with molecular dynamics is emerging as a successful approach to identify potential glutamate transporter isoforms with pharmacological potential [8] and identifying initial transporter conformations [14]. This study presents a comprehensive investigation of splice isoforms of the glutamate transporter EAA1, employing an integrative approach that combines multi-omics analysis, artificial intelligence-guided computational modeling, and 500ns full-atom molecular dynamics simulations. By systematically examining the structural and self-assembly characteristics of these truncated variants, we aim to understand the previously unrecognized structural adaptations that emerge from protein truncation and their potential significance in neural molecular processes.

We hereby report the discovery of two novel truncated isoforms of the glutamate transporter, which form stable β-sheet-enriched oligomers, representing a previously unobserved class of protein structures in glutamate transporters. Via identifying helix-to-sheet transformation dynamics, our research addresses a critical gap in the current understanding of glutamate transporter structural biology, offering novel insights into the complex relationship between protein truncation, structural reorganization, and potential functional implications. The findings not only contribute to fundamental protein science but also provide potential new perspectives on the molecular mechanisms of neurodegenerative conditions.

## Results and Discussions

### Comparative genomics analysis and the discovery of soluble truncated EAA1 isoforms

For the functional analysis of the splice variants, protein length was fundamental. This is because of the certain helices is essential for interaction with canonical EAA1, and isoforms required to contain these helices to interact with canonical EAA1 [8]. The analysis of isoforms derived from the canonical EAA1 protein (UniProt ID: P43003) previously yielded valuable insights into the physicochemical properties of truncated sequences [8]. These truncated variants have been shown to interfere with the canonical protein’s ability to form oligomers, which is a critical step in its functional activity within cellular membranes [8]. Surprisingly, certain isoforms were shown to produce self-assemblies, and some interactions exhibited a smaller interface area than those needed to inhibit canonical EAA1 [8]. On the other hand, previously mapped isoforms were focused on the 15–85% length of canonical transporter, with inhibitory effects on oligomerization [8]. For the analysis of smaller splice peptides, a total of nine isoforms were identified and curated for this study, focusing on those with lengths less than or equal to 15% of the canonical full-length structure.

The sampled isoforms were computationally mapped using UniProt’s gene-centric approach, which integrates data from Ensembl, EnsemblGenomes, model organism databases, and original sequencing projects [15]. The UniProt accession numbers for the nine isoforms under investigation were: A0A7P0T9P1, A0A7P0T8Q1, A0A7P0T7Q9, A0A7P0Z4F7, A0A7P0T8P5, A0A7P0TAF5, A0A7P0T8R2, A0A7P0T911, and A0A7P0T8H5. The physicochemical characterization of truncated EAA1 isoforms provided insights into their solubility and potential functional roles. The analysis revealed a diverse range of molecular weights and isoelectric points across the isoforms, reflecting the influence of truncation on protein properties (Table 1). These variations may have functional implications, particularly in the context of isoform-specific interactions, stability, and cellular localization [8]. In silico docking and molecular dynamics suggest that certain EAAT truncations have high binding energy when complexed with canonical transporters. For instance, we previously predicted that isoforms containing the TM2 helix exhibit especially strong interactions with the full-length transporter, effectively “competing” with native subunits [8]. In contrast, isoforms lacking those helices (e.g. mimicking TM5) tend to preferentially self-assemble, which may reduce their inhibitory impact [8].

**Table 1.** Sampled Isoforms.

| **UniProt ID** | **Length**<br>**(aa)** | **Molecular**<br>**Weight**<br>**(Da)** | **Isoelectric**<br>**Point (pI)<sup>1</sup>** | **Aliphatic**<br>**index<sup>2</sup>** | **Grand Average**<br>**of**<br>**Hydropathicity**<br>**(GRAVY)<sup>2</sup>** |
| --- | --- | --- | --- | --- | --- |
| **A0A7P0T9P1** | 34 | 4006.94 | 8.25 | 128.82 | 0.821 |
| **A0A7P0T8Q1** | 133 | 15278.09 | 9.88 | 89.32 | -0.078 |
| **A0A7P0T7Q9** | 71 | 8108.43 | 9.90 | 82.25 | -0.541 |
| **A0A7P0Z4F7** | 23 | 2683.39 | 9.51 | 140.00 | 0.430 |
| **A0A7P0T8P5** | 133 | 15368.47 | 10.06 | 101.73 | 0.249 |
| **A0A7P0TAF5** | 37 | 4277.05 | 11.10 | 50.00 | -1.208 |
| **A0A7P0T8R2** | 47 | 5264.02 | 7.16 | 29.15 | -0.713 |
| **A0A7P0T911** | 70 | 7921.41 | 10.17 | 75.14 | -0.366 |
| **A0A7P0T8H5** | 25 | 2799.33 | 6.73 | 31.20 | -0.336 |

<sup>1</sup>For each isoform, ExPASy web application (https://web.expasy.org/compute_pi/) was utilized to calculate the isoelectric points (pI) and molecular weights (MW) of the proteins.
<sup>2</sup>ProtParam, a computational tool, was employed to assess solubility profiles based on sequence-derived parameters.

Surprisingly, sequence analysis now reveals that some EAAT1 splice isoforms are highly hydrophilic and likely soluble. Table 1 shows multiple human *SLC1A3-*derived isoforms (e.g. UniProt A0A7P0TAF5, A0A7P0T8R2, A0A7P0T8H5) with grand average of hydropathicity (GRAVY) <0. This observation suggests that truncation impacts the hydrophobicity and folding patterns of these isoforms, potentially altering their interaction with lipid bilayers. These hydrophilic variants would not stably insert into lipid bilayers and so are expected to remain in aqueous environments. For instance, although some 133-residue isoforms (A0A7P0T8Q1, A0A7P0T8P5) have near-neutral GRAVY (–0.078 and +0.249), others (A0A7P0TAF5, A0A7P0T8R2) are markedly soluble. The 37-amino-acid isoform A0A7P0TAF5 has GRAVY = –1.208, indicating that it is a strongly hydrophilic peptide. The truncation therefore profoundly alters folding and membrane affinity: high aliphatic index or GRAVY of the short isoforms suggests the remaining sequence is dominated by polar residues. This implies that the splicing event effectively “trades” transmembrane segments for soluble regions.

The discovery of soluble isoform assemblies (Table 1), alongside the previously identified inhibitory potential of certain truncations, raises questions about their precise biological roles and the mechanisms underlying their functional divergence. The existence of water-soluble EAAT isoforms suggests new regulatory possibilities. Unlike membrane-bound truncations that inhibit via hetero-oligomerization [8], soluble fragments cannot assemble into the plasma membrane. Instead, they may act by different mechanisms. For example, a soluble isoform could bind to canonical EAATs in the ER lumen (if they carry luminal loops) or interact with cytosolic partners, thereby influencing transporter maturation or trafficking. They might also compete for binding proteins that normally associate with EAA1, indirectly modulating the amount of functional transporter. Alternatively, if secreted, they could act as short decoys or signaling molecules outside the cell (though none of the isoforms here has a clear signal peptide). In general, the change in hydrophobicity and isoelectric point means each isoform will have a distinct cellular localization. Previous studies have suggested that engineered soluble variants of glutamate transporters may have potential applications in diagnostics [16]. Truncated isoforms have long been recognized as valuable targets in the context of inhibitor design [17]. In particular, soluble truncated variants may represent an even more promising avenue for diagnostic applications, as their lack of membrane anchoring typically makes them more accessible for structural and functional characterization, thereby facilitating research and development. Moreover, this biochemical property often allows for easier purification, storage, and manipulation in experimental settings compared to their membranous counterparts [16]. Furthermore, we previously discovered that a natural soluble TLR5 variant in fish species could be evolutionary adapted for distinct signaling functions [18]. This TLR5 isoform appears to have undergone dynamic variational changes to serve distinct signaling roles, potentially environmentally diverging from the canonical pathogen-recognition pathway of the membrane-bound form [18]. At the very least, the presence of soluble isoforms of EAA1 raises questions on independent signaling functions.

### Structural modelling and self-assembly

To establish structural characterization and analyze self-binding of sampled isoforms (soluble and insoluble) we predict their multimeric structures. After the initial monomer predictions, hexametric models were generated for each isoform to evaluate their potential for beta-sheet formation. Among the predicted hexamers, isoforms A0A7P0TAF5, A0A7P0Z4F7, and A0A7P0T8H5 demonstrated the ability to form beta sheets, albeit with varying confidence scores as predicted by AlphaFold3. To further investigate the capacity for beta-sheet formation and the influence of assembly size, structural predictions were repeated for these three isoforms at various oligomeric states, including tetramers, pentamers, hexamers, heptamers, octamers, dodecamers, hexadecamers, and icosamers (4, 5, 6, 7, 8, 12, 16, and 20 copies, respectively). The confidence of the predicted structures was evaluated based on plDDT scores, with surface areas having plDDT > 70 being considered reliable.

Among the modeled structures, the octameric assembly of A0A7P0Z4F7 and the hexameric assembly of A0A7P0TAF5 emerged as the most confident predictions, exhibiting well-defined beta-sheet regions and favorable structural integrity (Supplementary Figure S1, Supplementary Figure S2). These models were characterized by high-confidence surface areas and consistent beta-sheet arrangements, suggesting their potential functional relevance. Conversely, predictions for A0A7P0T8H5 across all tested oligomeric states failed to achieve confidence thresholds indicative of reliable structural features. The absence of confident predictions for beta-sheet formation or stable assemblies led to the exclusion of A0A7P0T8H5 from further analysis. Interestingly, all sampled splice peptides were helical peptides in monomeric forms (Figure 1).

**Figure 1.**
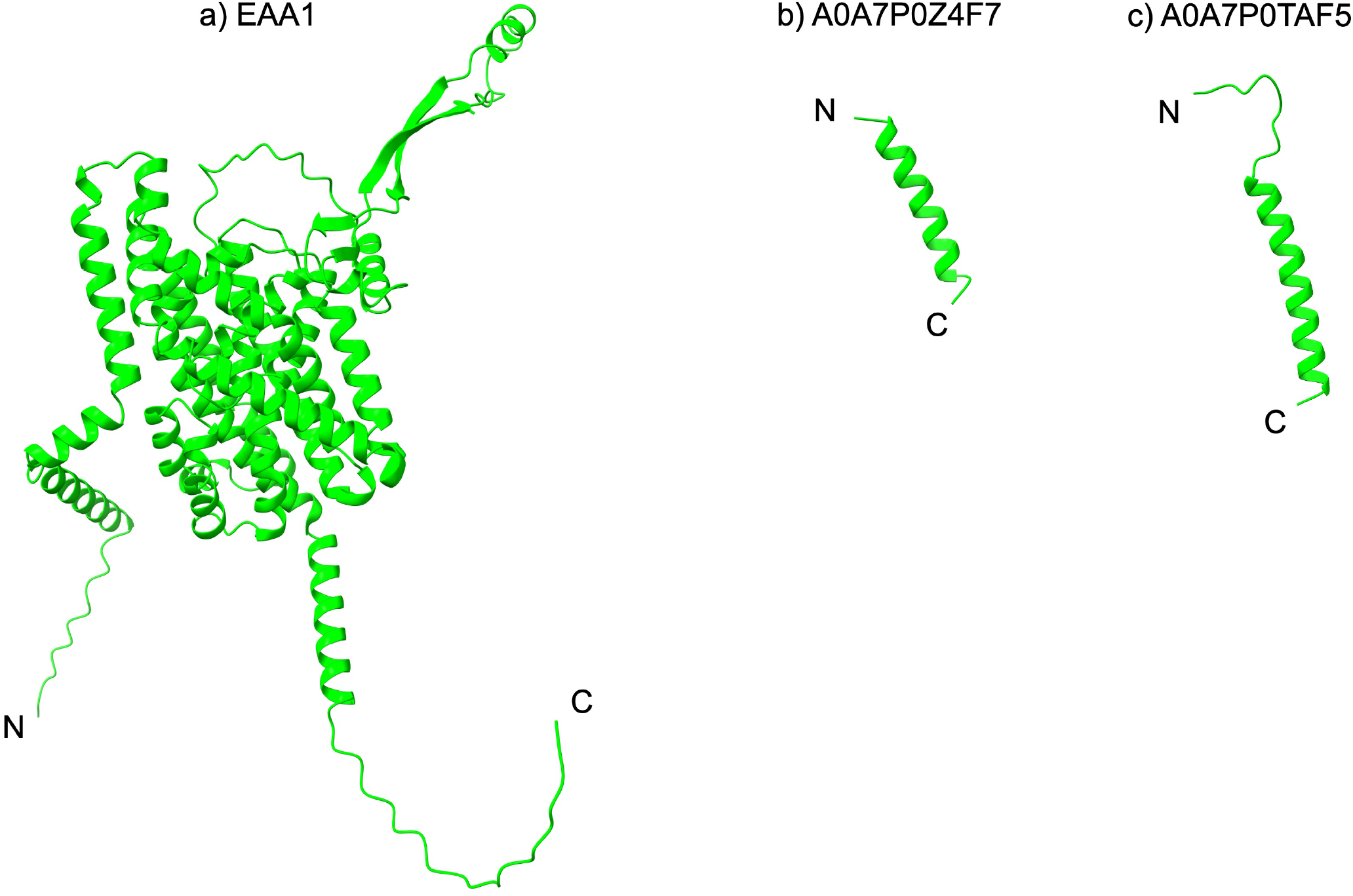
Structural representations of EAA1, A0A7P0Z4F7, and A0A7P0TAF5. **a**) The full-length structure of EAA1, a glutamate transporter, showcasing its characteristic transmembrane helices and extracellular loops, with N- and C-termini labeled. b) The truncated isoform A0A7P0Z4F7, represented as a single α-helix. c) The truncated isoform A0A7P0TAF5, also represented as a single α-helix, with slight differences in loop organization compared to A0A7P0Z4F7, with the N- and C-termini labeled.

Sheet-helical transformations often feature a specific charge distribution [11,12,13]. This is characterized by negatively charged residues clustering near the N-terminus and positively charged residues near the C-terminus, which balances the inherent dipole moment of an α-helix, thereby also stabilizing the helical conformation [11,12,13]. Thus, a defining characteristic for the transformation is negatively charged residues (like aspartic acid or glutamic acid) locating towards the N-terminus and a corresponding cluster of positively charged residues (like arginine or lysine) towards the C-terminus [11,12,13]. Interestingly, this aligns with the splice peptides analyzed in this study. The 0A7P0Z4F7 has a 5ASP residue, while A0A7P0TAF5 has a 2THR residue. The arrangement also allows for ionic complementarity and extensive ionic bonding on the hydrophilic face of the β-sheet assembly. The transition between these two stable states can be abrupt and often appears direct, without a detectable random-coil intermediate [12,13]. For example, heating a peptide above a threshold temperature triggers a rapid conversion from β-sheet to α-helix [12]. Conversely, the return from the α-helical state to the thermodynamically more stable β-sheet assembly is significantly slower, sometimes taking weeks, highlighting kinetic barriers and hysteresis in the process [12]. This phenomenon is sensitive to ionic strength, which tends to stabilize the β-sheet form and inhibit the transformation [11,12,13] The study of these sheet-helical transformations provides a valuable model system for understanding protein folding dynamics, protein-protein interactions, and the conformational changes associated with amyloid formation in neurodegenerative diseases [11,12,13].

### Analysis of Beta Sheet Dynamics of A0A7P0Z4F7 Octamer

Molecular dynamics (MD) simulations were conducted to evaluate the stability and initial dynamics of AlphaFold3-predicted structures for the self-assembly of splice peptides. This computational approach followed methodologies established in prior studies on glutamate transporters [8,14]. The predicted octameric assembly of A0A7P0Z4F7 was analyzed across a 500ns simulation window. Structural snapshots from the beginning (0ns) and end (500ns) of the simulation revealed significant conformational changes. At 0ns, the octamer displayed a tightly packed beta-sheet configuration (Figure 2a). By 500ns, a more dynamic and slightly twisted beta-sheet architecture emerged, reflecting structural relaxation and adaptive folding within the solvent environment (Figure 2b). These transitions are consistent with the expected flexibility of beta-sheet-rich oligomers in aqueous conditions. The twisting is accentuated by the opposing directional rotations of the beta strands (Supplementary Figure S3). The symmetry observed during bending underscores the organized nature of this structural transition, suggesting a cooperative effect among beta-strands. The twisting also facilitates tighter packing of beta-strands, which can enhance hydrophobic core interactions. Root Mean Square Fluctuation (RMSF) analysis over the 500ns trajectory revealed a periodic pattern of flexibility across the beta-sheet protein (Supplementary Figure S4). Regions corresponding to loop segments and termini exhibited higher atomic fluctuations (up to ~1.5 nm), whereas core beta strands maintained low RMSF values (~0.3–0.6 nm), indicating structural stability. This pattern supports the preservation of the beta-sheet scaffold throughout the simulation. The transition from a planar to twisted beta-sheet conformation may represent an early aggregation-prone state. Misfolding of EAAT1 isoforms could predispose them to aggregate into fibrillar structures, thereby potentially contributing to neurodegenerative conditions such as Alzheimer’s or ALS.

**Figure 2.**
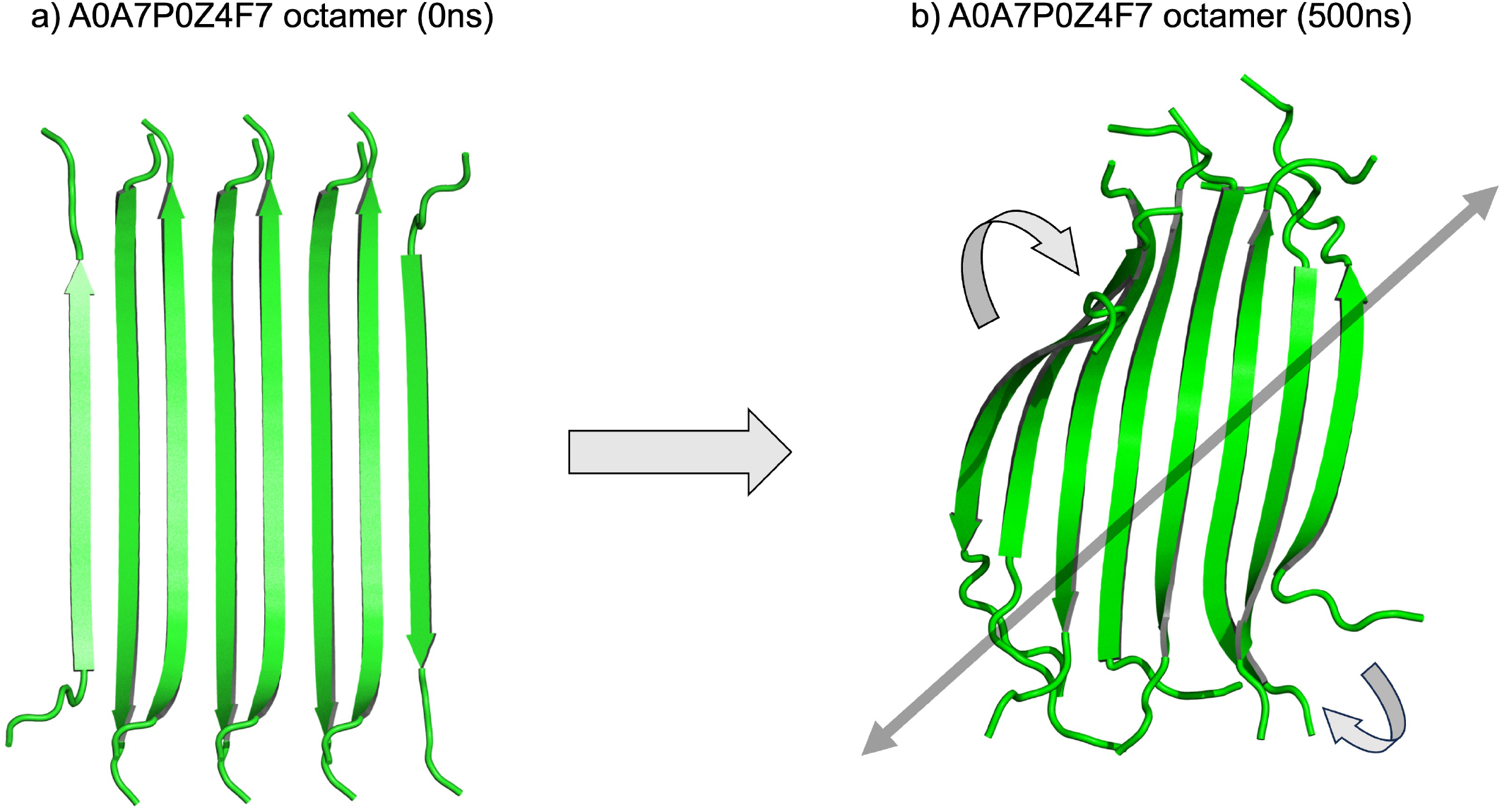
Structural evolution of the A0A7P0Z4F7 octamer during a 500ns molecular dynamics simulation. a) Initial configuration (0ns) of the A0A7P0Z4F7 octamer, displaying a planar, parallel arrangement of β-sheets. b) Final configuration after 500ns of simulation, showing a compact, twisted β-barrel-like structure with significant conformational changes, as indicated by the rotational movements (curved arrows) and diagonal axis alignment (gray arrow).

### Amphipathic Zonation and Structural Hierarchy

Further analysis focused on the distribution of hydrophobic and hydrophilic regions on the octameric surface (Figure 3). A0A7P0Z4F7 exhibited alternating hydrophobic and hydrophilic patches, with a symmetrical distribution along the longitudinal axis. This pattern suggests potential functional implications in membrane association and protein-protein interactions. Notably, hydrophobic regions aligned with the beta-sheet planes, while hydrophilic patches provided solvent accessibility, further stabilizing the oligomer in solution.

**Figure 3.**
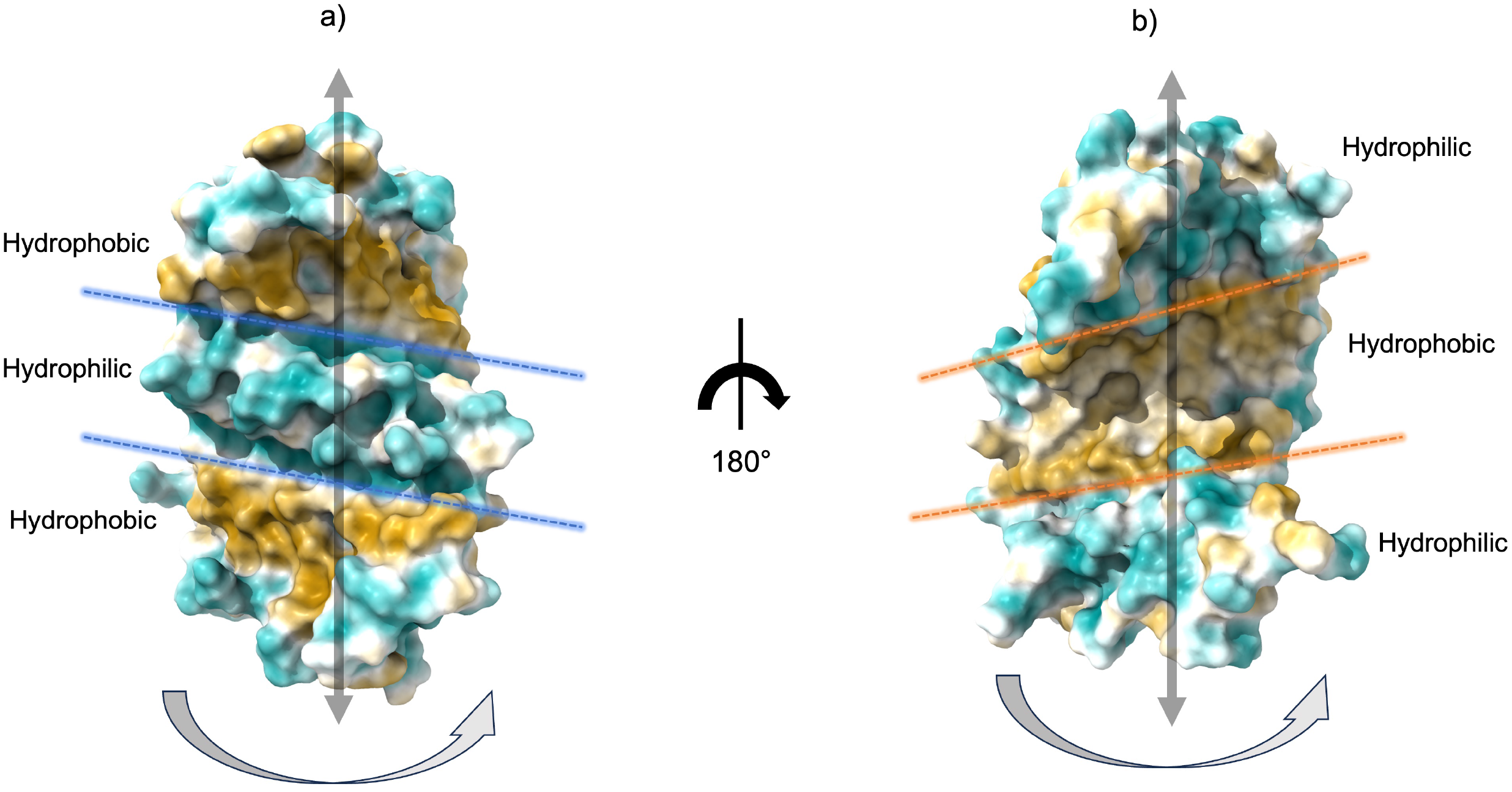
Final configuration of the A0A7P0Z4F7 octamer after 500ns of simulation. Surface representations are colored by hydrophobicity (hydrophobic regions in gold, hydrophilic regions in cyan). (a) Front view showing alternating hydrophobic and hydrophilic surface bands along the longitudinal axis. (b) Reverse view after 180° rotation, highlighting the complementary distribution of hydrophilic and hydrophobic regions. Blue and orange dashed lines indicate the approximate boundaries between hydrophobic and hydrophilic surface zones. Gray vertical arrows shows the longitudinal axis, and curved arrows indicate the viewing rotation between panels. The protein is in solution with Monte-Carlo placed K+ CL− ions (neutralizing, concentration[=[0.15 M) (Methods).

For A0A7P0TAF5, the initial conformation is characterized by a well-ordered, parallel beta-sheet arrangement. The transition from a flat to a twisted state could indicate intermediate states in either functional conformational changes or aggregation pathways. Twisted beta-sheets are often precursors to more compact oligomeric or fibrillar states [19,20]. Compared to the previous structure, this variant exhibits an increased amplitude of fluctuation at multiple sites, suggesting enhanced dynamic behavior possibly due to sequence variation or altered solvent exposure (Supplementary Figure S5). Accordingly, A0A7P0TAF5 has GRAVY = –1.208, while A0A7P0Z4F7 has a GRAVY score of 0.430 (Table 1). The RMSF profile over 500ns reveals a repeating fluctuation pattern with distinct peaks reaching up to ~2.0 nm, indicating pronounced flexibility in loop or solvent-exposed regions of the beta-sheet protein. In contrast, the central beta-sheet regions show consistently low fluctuations (~0.4–0.6 nm), reflecting high structural rigidity.

The tripartite distribution of hydrophilic-hydrophobic-hydrophilic surface character in A0A7P0Z4F7 suggests a membrane-inserting β-sheet oligomer (the isoform also has a positive GRAVY score). The central hydrophobic belt aligns with the lipid bilayer core, while the flanking hydrophilic faces remain solvent-exposed or interface with the polar head groups of lipid membranes. The vertical arrow through the central axis suggests the direction of membrane penetration or alignment, with the hydrophobic band acting as a membrane-spanning region. This is analogous to Aβ oligomers forming β-barrel pore structures [20]. Such amphipathic β-sheet assemblies may have broad implications such as neurotoxicity via Ca^2^□ dysregulation (e.g., Aβ or tau oligomers) and seeding and propagation in prion-like diseases [21,22,23]. On the other hand. In the rotated view, it is evident that the middle section of the β-sheet oligomer is predominantly hydrophilic, flanked by hydrophobic regions on either side (Figure 3). This inversion from the usual hydrophobic-core paradigm has important implications. This pattern may reflect a protofibrillar or β-sheet oligomer, capable of lateral association via hydrophobic faces and axial stacking via hydrogen bonding. A hydrophilic midsection in an oligomer could facilitate aqueous solvation, preventing premature aggregation until proper membrane insertion or cellular targeting occurs.

### β-Sheet Octamerization of the A0A7P0Z4F7 Induces Localized Upper Leaflet Membrane Pitting

To evaluate the membrane remodeling properties of the A0A7P0Z4F7 isoform of EAAT1, we conducted a 250ns all-atom molecular dynamics simulation of the octameric complex embedded in a lipid bilayer. Structural stabilization coincided with significant topological deformation of the surrounding membrane environment, particularly manifesting as upper bilayer pitting directly beneath the protein interface (Figure 4, Supplementary Figure S6). Quantitative geometrical analysis of membrane curvature deformation revealed a distinct pitting effect marked by vertical displacement (AD = 10.7 Å) from the bilayer midplane (line BC), with approximately orthogonal to the lipid headgroup axis (while ∠BAC = 120.6°). The vertices of the pitting triangle (A–B–C–D) were emphasizing a localized vertical depression beneath the octamer’s β-sheet-rich interaction core. The displacement vector AD bisects the B–C line, indicating a local curvature stress imposed by the isoform assembly. The quadrilateral CABD formed by the surrounding phosphate atoms further defines a localized distortion zone, effectively mapping the spatial domain of curvature influence imposed by the oligomer.

**Figure 4.**
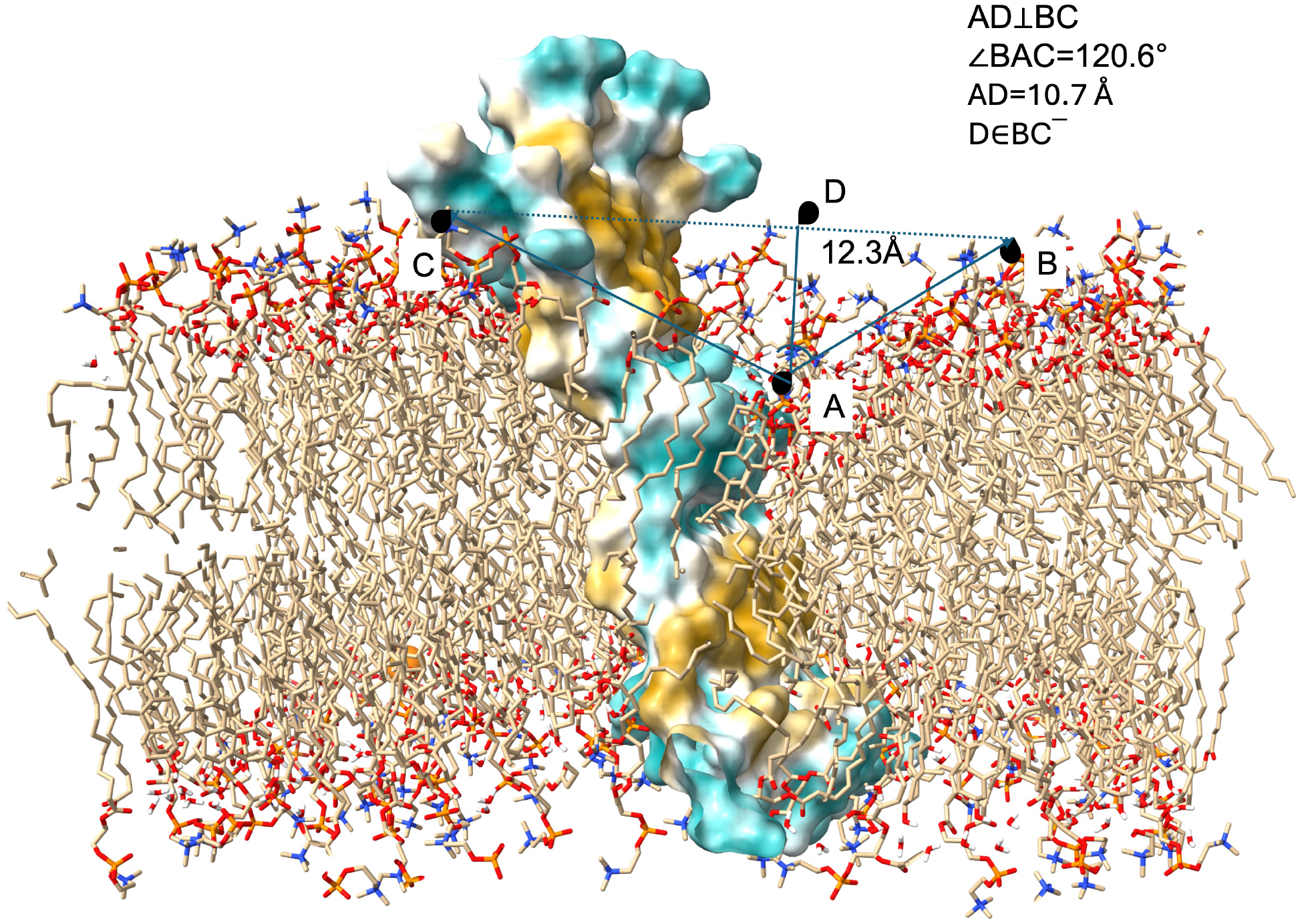
Upper layer membrane pitting via A0A7P0Z4F7 octamer after 250ns full-atom MD simulation in lipid bilayer. The protein surface is colored by hydrophobicity (hydrophobic regions in gold, hydrophilic regions in cyan). A, B, C is lipid head phosphates of POPC. Key reference points (A–D) and geometric parameters are indicated, with AD measured at 10.7 Å and ∠BAC = 120.6°. The initial orientation and depth of membrane insertion were refined using the PPM 2.0 method. The lipid bilayer composition consisted of 70% POPC and 30% cholesterol in both leaflets, consistent with our previous studies. See Supplementary Figure S6 for further information and measurements.

This β-sheet-mediated oligomerization-induced membrane perturbation diverges significantly from canonical EAAT1 assemblies or isoform dimers, which typically maintain membrane planarity or induce undulations [8]. This is further supported by the observation that the β-sheet interfaces are aligned laterally with the lipid headgroups, thereby enabling extended hydrogen bonding networks and possible interactions. Such β-sheet scaffolding could rigidify membrane-proximal regions, and in this case, likely induces local tension asymmetry across the bilayer, resulting in a concave bending toward the cytoplasmic side. The observed curvature strongly suggests that the A0A7P0Z4F7 isoform acts as a curvature scaffold, likely stabilizing via β-strand interactions at the membrane–protein interface (Supplementary Figure S7, Supplementary Figure S8). Additionally, the pitting effect could stabilize in tandem with the maturation of β-sheet contacts, as indicated by root mean square fluctuation (RMSF) analysis, which shows decreased mobility in residues participating in inter-protomer β-strand interactions (Supplementary Figure S4). These residues are topologically aligned with the inner face of the membrane depression, suggesting a mechanistic link between quaternary structure assembly and curvature induction.

The biological relevance of such membrane deformation may extend to transporter function and oligomer-specific regulation. For instance, localized pitting could create a permissive microenvironment for lipid sorting or cytosolic protein recruitment [24]. Alternatively, curvature-induced stress may signal membrane trafficking events or contribute to the partitioning of EAA1 isoforms into specialized membrane domains such as astrocytic processes, or lipid rafts. These findings highlight a potential mechanism by which helical EAA1 isoforms may engage in adaptive β-structural rearrangements to modulate local lipid architecture, contributing to higher-order spatial organization in synaptic membranes.

### Conformational Transformation of the A0A7P0TAF5 Hexamer

We further analyzed the second isoform (A0A7P0TAF5) that shows β-strand-rich structural predictions. The initial configuration of the A0A7P0TAF5 hexamer at 0ns displays a planar, stacked arrangement of six β-strand-rich monomers. These monomers are organized in a parallel β-sheet alignment, forming an extended fibrillar architecture (Figure 5). The inter-subunit interactions appear uniform and predominantly linear, suggesting a metastable oligomeric state reminiscent of early amyloidogenic intermediates. After 500ns of molecular dynamics simulation in water solvent, the hexamer adopts a markedly different conformation characterized by significant rotational and curvilinear reorganization of the β-strands. A 180° rotation along the vertical axis shows a barrel-like topology with distinct structural polarity (Figure 5). This transformation involves a global twisting motion, accompanied by an inward tilting of β-strands and reorientation of loop regions. The resulting structure exhibits features typical of β-barrel-like oligomers, including a potential hydrophilic core and peripheral exposure of hydrophobic residues. Interestingly, this is an unified feature of both sampled isoforms in this study, suggesting a common mechanistic explanation of isoform derived helix-to-sheet transformations of neurological transporter splice peptides.

**Figure 5.**
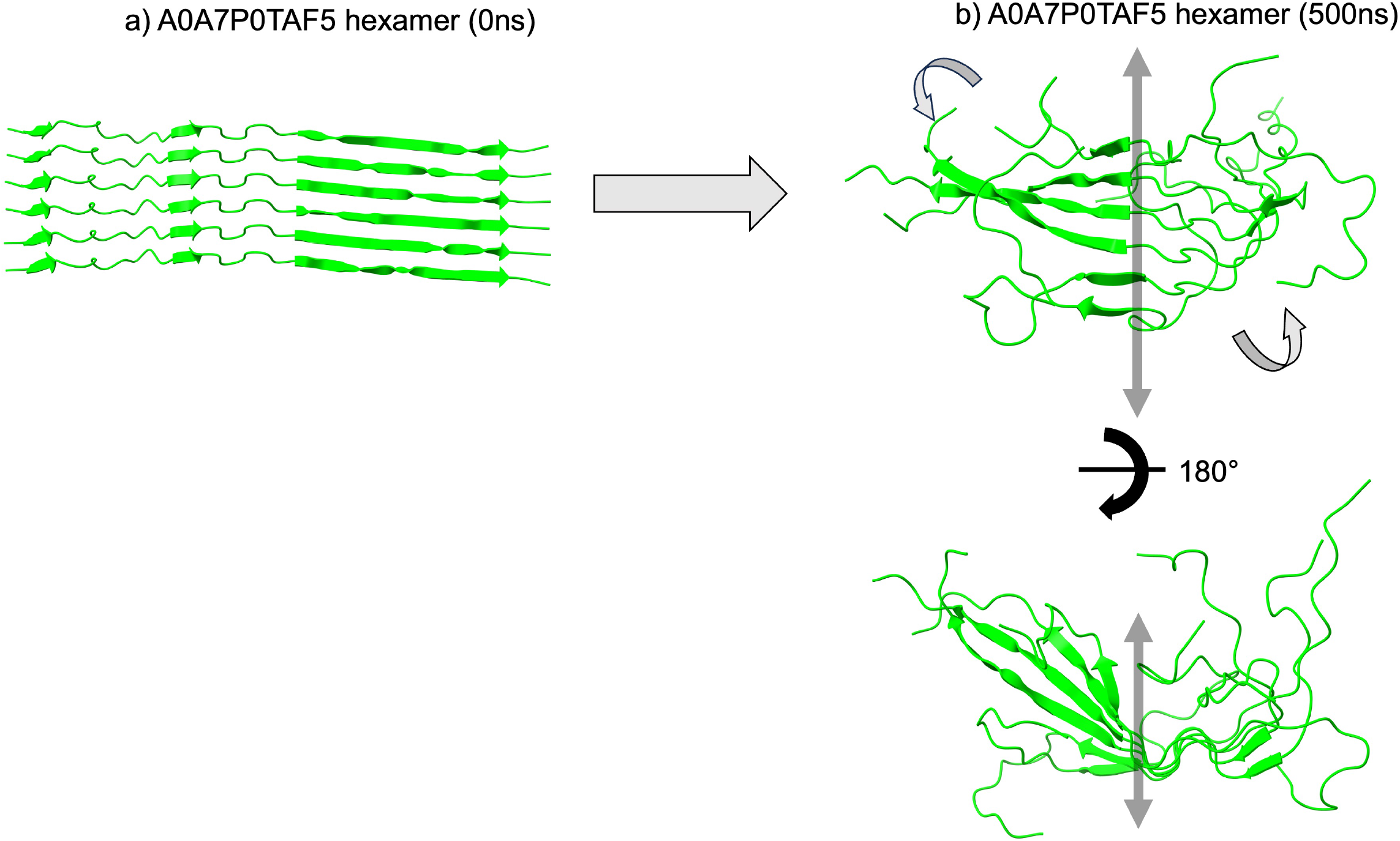
Conformational evolution of the A0A7P0TAF5 hexamer, showing structural changes in its beta-sheet arrangements over the course of a 500ns molecular dynamics (MD) simulation. a) Initial configuration (0ns) of the A0A7P0TAF5 hexamer, displaying a planar, parallel arrangement of β-sheets. b) Final configuration after 500ns of simulation. The protein is in solution with Monte-Carlo placed K+ CL− ions (neutralizing, concentration[=[0.15 M) (Methods).

## Discussion

In this study, we reveal two complementary layers of structural organization underlying the functional potential of helix-to-sheet oligomeric assemblies: the axial stratification of hydrophobic and hydrophilic surface regions, and the dynamic transition of β-sheet-rich oligomers into compact β-barrel architectures. These insights illuminate how molecular topology and surface chemistry co-evolve during oligomer assembly of glutamate transporters, with implications for both physiological and pathological processes. These findings suggest that both static surface hydrophobicity patterns and dynamic shape transitions determine the functional capacity of oligomeric assemblies. The co-existence of spatially ordered hydrophobic/hydrophilic domains and a propensity for β-barrel formation supports a model in which these oligomers may participate in membrane insertion, transient pore formation, or signaling through lipid interactions. Moreover, the hydrophilic central band observed in the static structure may facilitate initial aqueous solubility, with subsequent β-barrel closure enabling membrane partitioning, a behavior consistent with a metastable, conditionally active oligomeric intermediate.

These results not only advance our understanding of sequence-dependent oligomer morphologies but also provide a conceptual framework for interpreting the dual membrane-compatible and aggregation-prone nature of many β-rich oligomers implicated in disease. Future studies integrating mutational scans, membrane mimetic environments, and experimental validation (e.g., cryo-EM or patch-clamp electrophysiology) will be critical to confirming the functional roles of these potential metastable intermediates.

## Methods

### Protein sequence alignments and other characteristics

The protein sequence data were retrieved in FASTA format from the UniProt website (https://www.uniprot.org) [15]. The UniProt accession numbers for the canonical EAA1 is P43003. The corresponding isoforms were also fetched utilizing UniProt which identifies potential isoforms via automatic gene-centric mapping from eukaryotic reference proteomes. This mapping is based on gene identifiers from Ensembl, EnsemblGenomes, model organism databases and original sequencing projects [15]. The retrieved sequences were filtered to generate a dataset containing exclusively computationally mapped truncated isoforms. This study specifically analyses the isoforms of the length of %15 or less of the canonical full-length structures. Uniprot IDs for each isoform (count=9) were: A0A7P0T9P1, A0A7P0T8Q1, A0A7P0T7Q9, A0A7P0Z4F7, A0A7P0T8P5, A0A7P0TAF5, A0A7P0T8R2, A0A7P0T911, A0A7P0T8H5.

For each isoform, Expasy web application (https://web.expasy.org/compute_pi/) was utilized to calculate the isoelectric points (pI) and molecular weights (MW) of the proteins [25,26]. ProtParam, a computational tool, was employed to assess solubility profiles based on sequence-derived parameters [26].

### Comparative structural analysis

AlphaFold3 Program [27] was used for the structure predictions of the studied isoforms following the instructions at the website. After monomer predictions, hexamer structures were modelled for each isoform to assess overall ability to form beta sheets. Predicted hexamers of A0A7P0TAF5, A0A7P0Z4F7, and A0A7P0T8H5 formed beta sheets with differing confidence scores. On the other hand, the models could form beta sheets with different assemblies and copy counts. We repeated predictions for these 3 potentially beta sheet forming isoforms for 4,5,6,7,8,12,16 and 20 copies. The structures analyses according to confidently predicted surface areas (plDDT > 70). Among them, octamer of A0A7P0Z4F7 and hexamer of A0A7P0TAF5 were most confident predictions. A0A7P0T8H5 did not have any confident structures, thereby rejected from the sample.

### Molecular Dynamics Simulations

Molecular dynamics simulations were carried out on AlphaFold3-predicted structure variants in solution and membrane, following the approach used in our previous study on glutamate transporter interactions [8,14]. All simulations employed GROMACS version 2024.3 [28] and were run on Google Colab utilizing L4 GPUs, 106 GB of RAM, and 43 GB of VRAM. To maximize computational efficiency, simulations were parallelized across multiple cores within the virtual machine. Additionally, the GROMACS source code was recompiled to leverage all available cores with CUDA GPU acceleration enabled (DGMX_GPU=CUDA) and OpenMP threading (GMX_OPENMP_MAX_THREADS=128), specifically optimized for L4 GPUs as outlined in our recent benchmarking work [29]. Configuration files and simulation scripts, along with detailed step-by-step instructions, have been made publicly accessible [29].

Accordingly, solution-protein systems were built using the CHARMM-GUI web server [28,30,31,32]. The rectangle-shaped water box of edge distance 10Å was formed and solvated using explicit solvation. System pH for both membrane and water solvent systems were set to 7.0. The systems were built with neutralizing K^+^, Cl^−^ ions (concentration = 0.15M), which was determined through 2000 step Monte Carlo (MC) simulations using a primitive model. For membrane interactions of A0A7P0Z4F7 octamer, membrane– protein systems were constructed using the CHARMM-GUI Membrane Builder [33]. Proteins were placed in rectangular simulation boxes, and their protonation states were assigned based on the system’s local pH. The orientation and depth of membrane insertion were refined using the PPM 2.0 method, which takes into account hydrogen bonding patterns and the anisotropic nature of the water–lipid environment [34]. Lipid bilayers consisted of 70% 1-palmitoyl-2-oleoyl-sn-glycero-3-phosphocholine (POPC) and 30% cholesterol in both leaflets, consistent with our previous studies [8]. This composition was selected to reflect the key structural and functional features of a typical plasma membrane, in line with prior experimental and computational models [8].

All systems were solvated using the TIP3P water model, and simulations were conducted with the all-atom CHARMM36m force field [35]. To prepare the systems, a multi-step minimization and equilibration protocol, adapted from our previous work [8,14], was applied to both protein–membrane and protein–solution environments. Energy minimization was performed using the steepest descent algorithm, ensuring that the maximum force on any atom dropped below 1000 kJ/mol/nm. Subsequently, a 125ps equilibration phase was carried out for each system, following the standard CHARMM-GUI workflow [32], including NVT and NPT ensembles. Production molecular dynamics (MD) simulations were then executed for 500ns and 250ns, with trajectory snapshots saved every 0.5ns. Electrostatic interactions were computed using the Particle Mesh Ewald (PME), and both Coulombic and van der Waals interactions were truncated at 1.2 nm. The temperature was maintained at 303.15K using the Nose–Hoover thermostat, and pressure was controlled at 1 bar via the Parrinello–Rahman barostat with semi-isotropic coupling.

### Trajectory analysis

Comparative trajectory analyses were performed to assess the structural stability and dynamics of the systems. Equilibrated trajectories from the 500ns simulations were merged using the gmx traj module in GROMACS. The radius of gyration for each system was computed with the gmx gyrate tool to evaluate overall compactness. To quantify local flexibility, residue-level root mean square fluctuations (RMSF) of the Cα atoms were calculated over the full simulation period. The solvent-accessible surface area (SASA) of side chains was determined using a probe radius of 1.4Å, following standard protocols [36]. All plots derived from these analyses were generated and visualized using Grace (https://plasma-gate.weizmann.ac.il/Grace/). Structural snapshots and dynamic trajectories were examined using UCSF ChimeraX (version 1.8). Additionally, residue-wise root mean square deviations (RMSD) were computed and visualized for both membrane-bound and water-soluble systems using ChimeraX, enabling direct comparisons of conformational stability in different environments.

### Membrane Pitting Analysis

The membrane surface topography was quantified from an equilibrated membrane–protein simulation of A0A7P0Z4F7 octamer (250ns) using VMD (Visual Molecular Dynamics version 2.0) without the use of external plugins or extensions [37]. The membrane phosphate atoms (POPC) were selected to define the bilayer surface. The system was divided into a regular 10 × 10 grid along the x–y plane using the atomselect and measure minmax commands in the VMD Tcl console to obtain coordinate bounds. For each grid cell (i, j), the average z-coordinate of phosphate atoms within the cell was calculated by iterating over frames to produce a height map of the bilayer leaflet. Missing values (NaN) were assigned where no atoms were present in a given grid cell. The resulting height map was exported as a .dat file containing the i, j indices and the corresponding average z-coordinate values. Pits were identified as local depressions in the height profile, defined as contiguous grid cells with z-coordinates at least Δz nm below the local mean surface level. Depth and area of pits were computed by subtracting pit-region z values from the surrounding baseline and counting the number of grid cells per depression. A binary mask was generated for pit regions, and connected components were identified using scipy.ndimage.label. For each pit, the total area (number of bins × bin area), maximum depth (μ − minimum z), minimum z, mean z, and bin count were calculated. The results were stored in a DataFrame for sorting and further analysis. Visualization was performed via Matplotlib 3.9.0 by plotting the z-height map with overlaid red contours marking pit boundaries, enabling both qualitative and quantitative characterization of membrane depressions.

For the measurement of phosphate distances and geometric construction, membrane phosphate positions were identified and measured using PyMOL v.3.03 (Schrödinger, LLC). Following system preparation, equilibrium and 250ns production simulations; phosphate atoms from lipid headgroups were selected based on residue names corresponding to the bilayer composition (POPC). Specific phosphate atoms were chosen from the upper and lower leaflets to define geometric reference points (A, B, C, and D). A geometric triangle was constructed from the selected phosphates using PyMOL’s measurement objects, with distances and angles calculated for precise numerical reporting. Interatomic distances (AB, AC, BC) and the angle ∠BAC were obtained using PyMOL’s functions, while perpendicular distances (e.g., AD ⍰ BC) were determined by trigonometric calculations. This procedure allowed reproducible determination of lipid headgroup spatial relationships in the membrane–protein complex representation.

## Supplementary Information (SI)

Supplementary Information.pdf

Supplementary data file containing more detailed analyses on AlphaFold3 predictions, subunit interactions and membrane profiling, statistical analyses, and individual graphs supporting the main findings presented in the manuscript.

## Ethics Approval

Ethics approval was not required for this computational study as it did not involve animal subjects, human participants, and identifiable data.

## Consent to participate

Not applicable. This computational study did not involve human participants.

## Consent for publication

Not applicable. This computational study did not involve human participants.

## Availability of data and materials

Each statistical and computational analysis of this study, included with step-by-step instructions where possible, are publicly available to ensure repeatability. For more detailed information on the statistical analyses, input files and detailed outputs, including the AlphaFold calculations and codes to regenerate analyses, please visit the website: https://github.com/karagol-alper/glutamate-soluble-isoforms.

## Competing financial interests

None.

## Funding

The author(s) received no specific funding for this work.

## Acknowledgements

We thank Dr. Shuguang Zhang from Lab of Molecular Architecture, MIT for his inputs on how to present our data.

## Notes

https://github.com/karagol-alper/glutamate-soluble-isoforms

### Competing Interest Statement

The authors have declared no competing interest.

## References

1) Zhang, D., Hua, Z., & Li, Z. (2024). The role of glutamate and glutamine metabolism and related transporters in nerve cells. CNS neuroscience & therapeutics, 30(2), e14617. 10.1111/cns.14617

2) Lewerenz, J., & Maher, P. (2015). Chronic glutamate toxicity in neurodegenerative diseases—what is the evidence?. Frontiers in neuroscience, 9, 170294. 10.3389/fnins.2015.00469

3) O’Donovan, S. M., Sullivan, C. R., & McCullumsmith, R. E. (2017). The role of glutamate transporters in the pathophysiology of neuropsychiatric disorders. NPJ schizophrenia, 3(1), 32. 10.1038/s41537-017-0037-1

4) Parkin, G. M., Udawela, M., Gibbons, A., & Dean, B. (2018). Glutamate transporters, EAAT1 and EAAT2, are potentially important in the pathophysiology and treatment of schizophrenia and affective disorders. World journal of psychiatry, 8(2), 51. 10.5498/wjp.v8.i2.51

5) Walsh, T., McClellan, J. M., McCarthy, S. E., Addington, A. M., Pierce, S. B., Cooper, G. M., … & Sebat, J. (2008). Rare structural variants disrupt multiple genes in neurodevelopmental pathways in schizophrenia. science, 320(5875), 539–543. 10.1126/science.1155174

6) Liu, Q., Fang, L., & Wu, C. (2022). Alternative splicing and isoforms: from mechanisms to diseases. Genes, 13(3), 401.

7) O’Donovan, S. M., Hasselfeld, K., Bauer, D., Simmons, M., Roussos, P., Haroutunian, V., Meador-Woodruff, J. H., & McCullumsmith, R. E. (2015). Glutamate transporter splice variant expression in an enriched pyramidal cell population in schizophrenia. Translational psychiatry, 5(6), e579. 10.1038/tp.2015.74

8) Karagöl, A., Karagöl, T., Li, M., & Zhang, S. (2024). Inhibitory Potential of the Truncated Isoforms on Glutamate Transporter Oligomerization Identified by Computational Analysis of Gene-Centric Isoform Maps. Pharmaceutical research, 10.1007/s11095-024-03786-z. Advance online publication. 10.1007/s11095-024-03786-z

9) Chiti, F., & Dobson, C. M. (2006). Protein misfolding, functional amyloid, and human disease. Annual review of biochemistry, 75, 333–366. 10.1146/annurev.biochem.75.101304.123901

10) Danielsson, J., Jarvet, J., Damberg, P., & Gräslund, A. (2005). The Alzheimer beta-peptide shows temperature-dependent transitions between left-handed 3-helix, beta-strand and random coil secondary structures. The FEBS journal, 272(15), 3938–3949. 10.1111/j.1742-4658.2005.04812.x

11) Altman, M., Lee, P., Rich, A., & Zhang, S. (2000). Conformational behavior of ionic self-complementary peptides. Protein science : a publication of the Protein Society, 9(6), 1095–1105. 10.1110/ps.9.6.1095

12) Zhang, S., & Rich, A. (1997). Direct conversion of an oligopeptide from a beta-sheet to an alpha-helix: a model for amyloid formation. Proceedings of the National Academy of Sciences of the United States of America, 94(1), 23–28. 10.1073/pnas.94.1.23

13) Zhang, S., & Altman, M. (1999). Peptide self-assembly in functional polymer science and engineering. Reactive and Functional Polymers, 41(1-3), 91–102. 10.1016/S1381-5148(99)00031-0

14) Karagöl, A., Karagöl, T., & Zhang, S. (2024). Molecular Dynamic Simulations Reveal that Water-Soluble QTY-Variants of Glutamate Transporters EAA1, EAA2 and EAA3 Retain the Conformational Characteristics of Native Transporters. Pharmaceutical research, 41(10), 1965–1977. 10.1007/s11095-024-03769-0

15) UniProt Consortium, T. (2018). UniProt: the universal protein knowledgebase. Nucleic acids research, 46(5), 2699–2699. 10.1093/nar/gkae1010

16) Karagöl, A., Karagöl, T., Smorodina, E., & Zhang, S. (2024). Structural bioinformatics studies of glutamate transporters and their AlphaFold2 predicted water-soluble QTY variants and uncovering the natural mutations of L-> Q, I-> T, F-> Y and Q-> L, T-> I and Y-> F. Plos one, 19(4), e0289644. 10.1371/journal.pone.0289644

17) Qing, R., Tao, F., Chatterjee, P., Yang, G., Han, Q., Chung, H., … & Zhang, S. (2020). Non-full-length water-soluble CXCR4QTY and CCR5QTY chemokine receptors: Implication for overlooked truncated but functional membrane receptors. Iscience, 23(12). 10.1016/j.isci.2020.101670

18) Karagöl, A., & Karagöl, T. (2025). Adaptation to Solvent Environment in Toll-like Receptor 5: A Comparative Evolutionary Analysis of Membrane-bound and Soluble Forms in Epinephelus coioides. bioRxiv, 2025–02. 10.1101/2025.02.28.640895

19) Kahler, A., Sticht, H., & Horn, A. H. (2013). Conformational stability of fibrillar amyloid-beta oligomers via protofilament pair formation–a systematic computational study. PloS one, 8(7), e70521. 10.1371/journal.pone.0070521

20) Diociaiuti, M., Bonanni, R., Cariati, I., Frank, C., & D’Arcangelo, G. (2021). Amyloid prefibrillar oligomers: the surprising commonalities in their structure and activity. International journal of molecular sciences, 22(12), 6435. 10.3390/ijms22126435

21) Malchiodi-Albedi, F., Paradisi, S., Matteucci, A., Frank, C., & Diociaiuti, M. (2011). Amyloid oligomer neurotoxicity, calcium dysregulation, and lipid rafts. International journal of Alzheimer’s disease, 2011(1), 906964. 10.4061/2011/906964

22) Wells, C., Brennan, S. E., Keon, M., & Saksena, N. K. (2019). Prionoid proteins in the pathogenesis of neurodegenerative diseases. Frontiers in molecular neuroscience, 12, 271. 10.3390/ijms22126435

23) Christensen, C. S., Wang, S., Li, W., Yu, D., & Li, H. J. (2024). Structural variations of prions and prion-like proteins associated with neurodegeneration. Current Issues in Molecular Biology, 46(7), 6423–6439. 10.3390/cimb46070384

24) McMahon, H. T., & Gallop, J. L. (2005). Membrane curvature and mechanisms of dynamic cell membrane remodelling. Nature, 438(7068), 590–596. 10.1038/nature04396

25) Bjellqvist, B., Basse, B., Olsen, E., & Celis, J. E. (1994). Reference points for comparisons of two□dimensional maps of proteins from different human cell types defined in a pH scale where isoelectric points correlate with polypeptide compositions. Electrophoresis, 15(1), 529–539. 10.1002/elps.1150150171

26) Gasteiger, E., Hoogland, C., Gattiker, A., Duvaud, S. E., Wilkins, M. R., Appel, R. D., & Bairoch, A. (2005). Protein identification and analysis tools on the ExPASy server (pp. 571–607). Humana press.

27) Abramson J, Adler J, Dunger J, et al. Accurate structure prediction of biomolecular interactions with AlphaFold 3. Nature 2024:1–3.

28) Abraham, M. J., Murtola, T., Schulz, R., Páll, S., Smith, J. C., Hess, B., & Lindahl, E. (2015). GROMACS: High performance molecular simulations through multi-level parallelism from laptops to supercomputers. SoftwareX, 1, 19–25. 10.1016/j.softx.2015.06.001

29) Karagöl, T., & Karagöl, A. (2024). Benchmarking GROMACS on Optimized Colab Processors and the Flexibility of Cloud Computing for Molecular Dynamics. bioRxiv, 2024-11. 10.1101/2024.11.14.623563

30) Kimura, M. (1979). Model of effectively neutral mutations in which selective constraint is incorporated. Proceedings of the National Academy of Sciences, 76(7), 3440–3444. 10.1073/pnas.76.7.3440

31) Wu, E. L., Cheng, X., Jo, S., Rui, H., Song, K. C., Dávila-Contreras, E. M., Qi, Y., Lee, J., Monje-Galvan, V., Venable, R. M., Klauda, J. B., & Im, W. (2014). CHARMM-GUI Membrane Builder toward realistic biological membrane simulations. Journal of computational chemistry, 35(27), 1997– 2004. 10.1002/jcc.23702

32) Jo, S., Kim, T., Iyer, V. G., & Im, W. (2008). CHARMM-GUI: a web-based graphical user interface for CHARMM. Journal of computational chemistry, 29(11), 1859–1865. 10.1002/jcc.20945

33) Jo, S., Lim, J. B., Klauda, J. B., & Im, W. (2009). CHARMM-GUI Membrane Builder for mixed bilayers and its application to yeast membranes. Biophysical journal, 97(1), 50–58. 10.1016/j.bpj.2009.04.013

34) Lomize, M. A., Pogozheva, I. D., Joo, H., Mosberg, H. I., & Lomize, A. L. (2012). OPM database and PPM web server: resources for positioning of proteins in membranes. Nucleic acids research, 40(Database issue), D370–D376. 10.1093/nar/gkr703

35) Huang, J., Rauscher, S., Nawrocki, G., Ran, T., Feig, M., de Groot, B. L., Grubmüller, H., & MacKerell, A. D., Jr (2017). CHARMM36m: an improved force field for folded and intrinsically disordered proteins. Nature methods, 14(1), 71–73. 10.1038/nmeth.4067

36) Eisenhaber, F., Lijnzaad, P., Argos, P., Sander, C., & Scharf, M. (1995). The double cubic lattice method: Efficient approaches to numerical integration of surface area and volume and to dot surface contouring of molecular assemblies. Journal of computational chemistry, 16(3), 273–284. 10.1002/jcc.540160303

37) Humphrey, W., Dalke, A., & Schulten, K. (1996). VMD: visual molecular dynamics. Journal of molecular graphics, 14(1), 33–38. 10.1016/0263-7855(96)00018-5

