## Supplementary Files for "Helix-to-Beta-Sheet Transition Drives Self-Assembly of Glutamate Transporter EAA1 Splice Peptides"

### Figures and Figure Legends

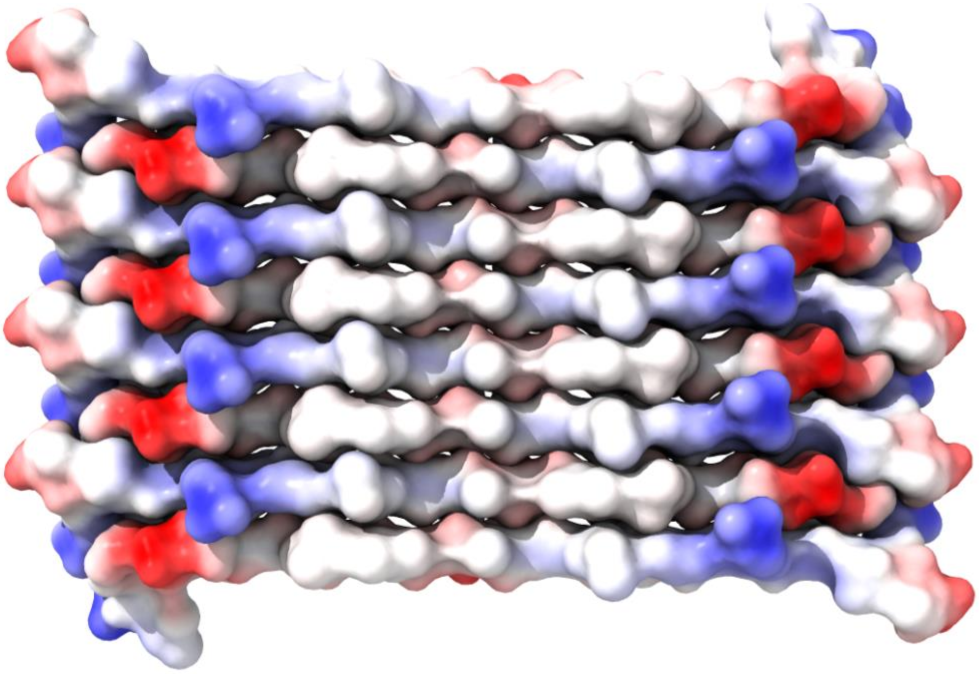

**Supplementary Figure S1. Predicted Octameric Assembly of A0A7P0Z4F7 Protein from AlphaFold3.** The surface representation of the AlphaFold3-predicted octameric structure of the uncharacterized protein A0A7P0Z4F7 is shown. The model reveals a highly ordered, stacked β-sheet-like architecture arranged in a fibrillar fashion. The electrostatic surface is colored by charge: positive (blue), negative (red), and neutral (white), highlighting alternating polarity and potential charge complementarity at subunit interfaces. This arrangement suggests a potential amyloid-like or filamentous assembly, possibly indicative of functional or pathological aggregation behavior.

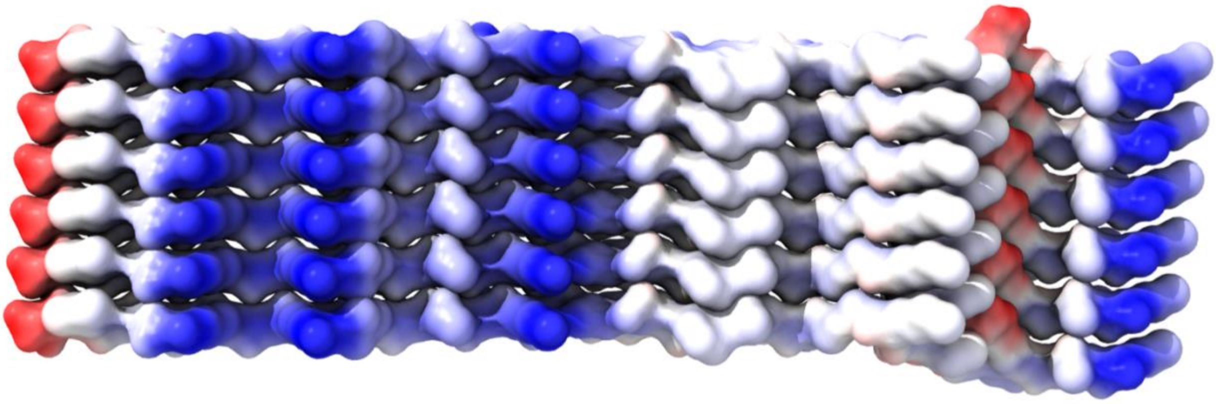

**Supplementary Figure S2. Predicted Assembly of A0A7P0TAF5 Hexamer from AlphaFold3.** The surface representation of the AlphaFold3-predicted octameric structure of the uncharacterized protein A0A7P0TAF5 Hexamer is shown. The model reveals a highly ordered, stacked β-sheet-like architecture arranged in a fibrillar fashion. The electrostatic surface is colored by charge: positive (blue), negative (red), and neutral (white), highlighting alternating polarity and potential charge complementarity at subunit interfaces. This arrangement suggests a potential amyloid-like or filamentous assembly, possibly indicative of functional or pathological aggregation behavior.

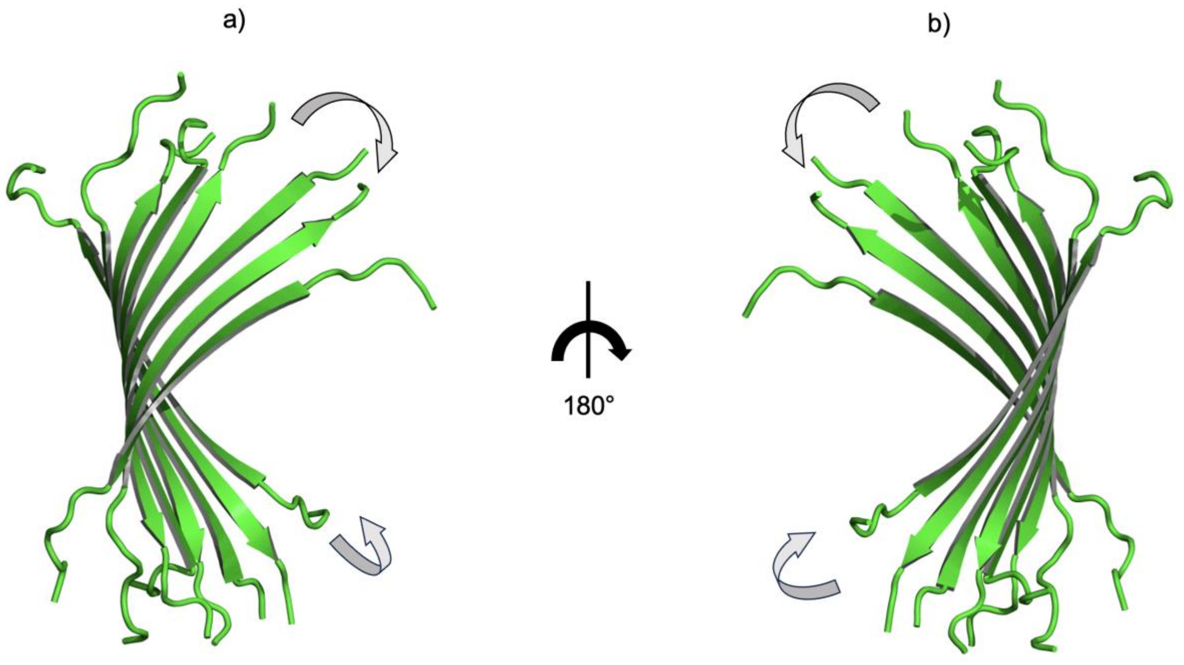


**Supplementary Figure S3. Final configuration of the A0A7P0Z4F7 octamer after 500 ns of simulation.** a, b) A compact, twisted β-barrel-like structure with significant conformational changes, as indicated by the rotational movements (curved arrows)

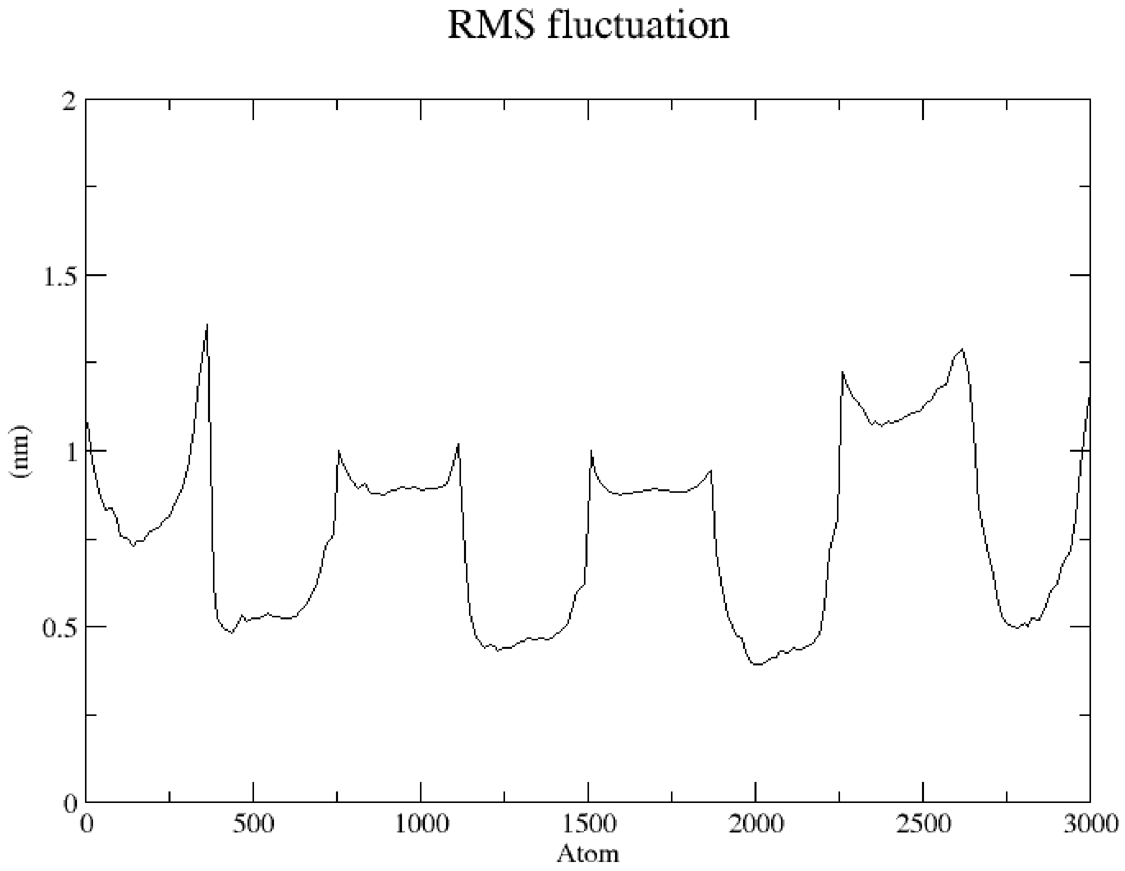


**Supplementary Figure S4. RMSF profiles during 500ns full-atom molecular dynamic simulations of A0A7P0Z4F7 octamer in water solvent.**

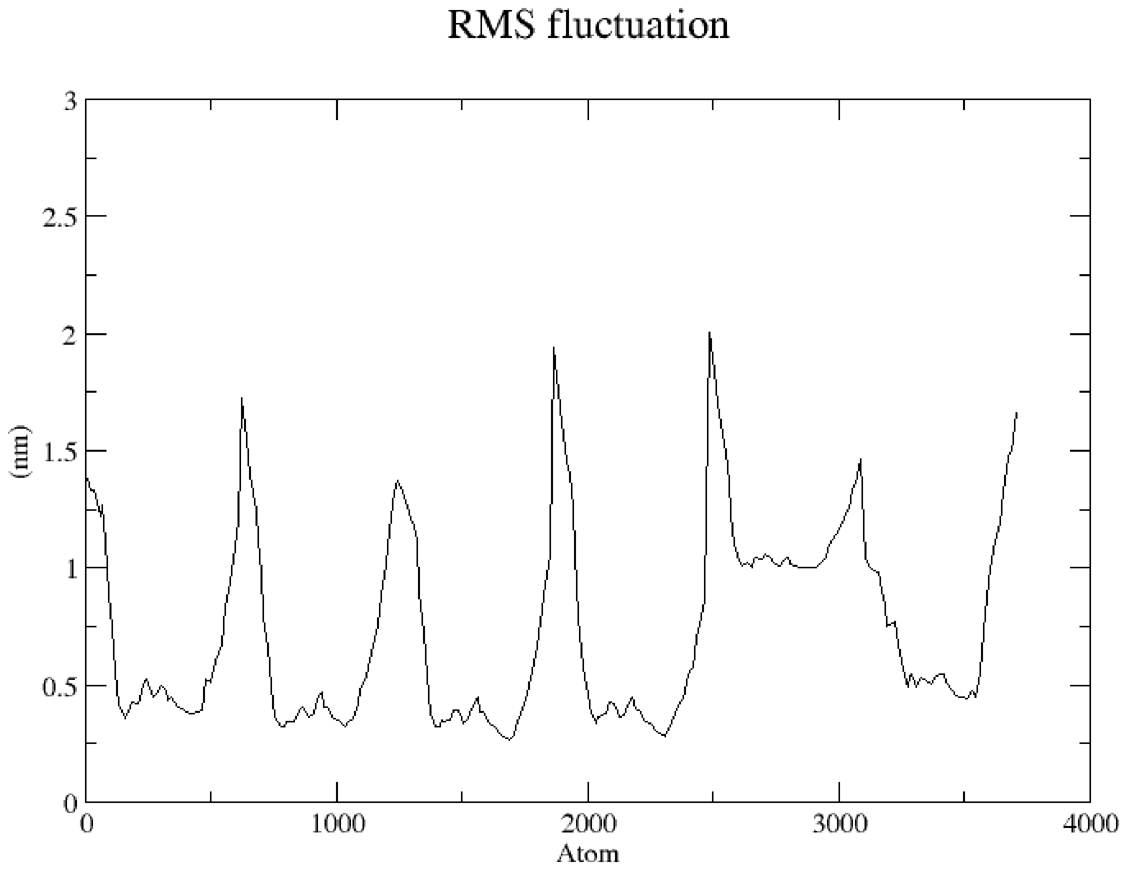


**Supplementary Figure S5. RMSF profiles during 500ns full-atom molecular dynamic simulations of A0A7P0TAF5 Hexamer in water solvent.**

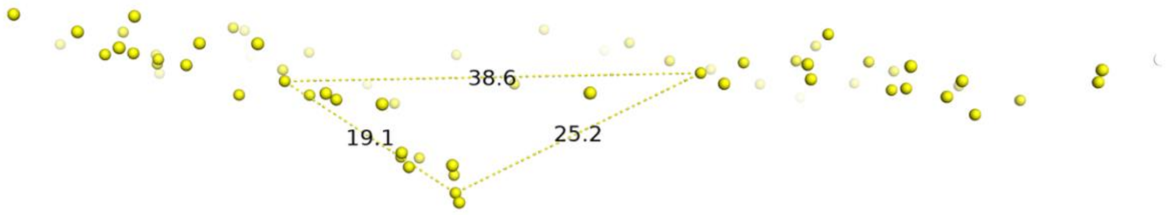


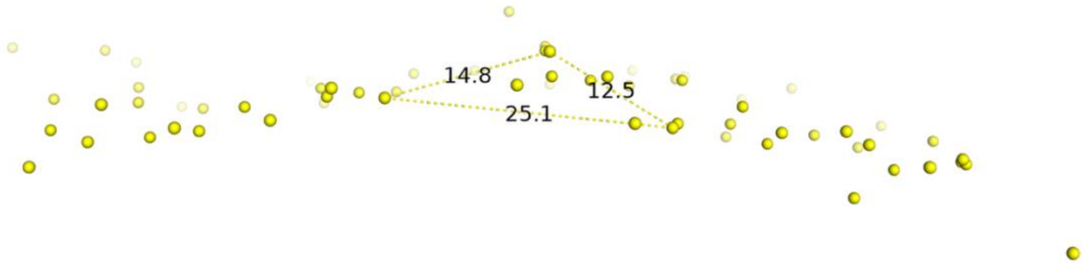


**Supplementary Figure S6. Lipid head phosphates of the membrane-protein system after 250ns full-atom MD simulation.** Side a = 38.6. Side b = 19.1. Side c = 25.2. Angle ∠A = 120.606°. Angle ∠B = 25.207°. Angle ∠C = 34.187°. Area = 207.1334. Perimeter p = 82.9. Semiperimeter s = 41.45. Height ha = 10.7323. Height hb = 21.68936. Height hc = 16.43916. Median ma = 11.28871. Median mb = 31.16565. Median mc = 27.72409. Inradius r = 4.99719. Circumradius R = 22.4239.

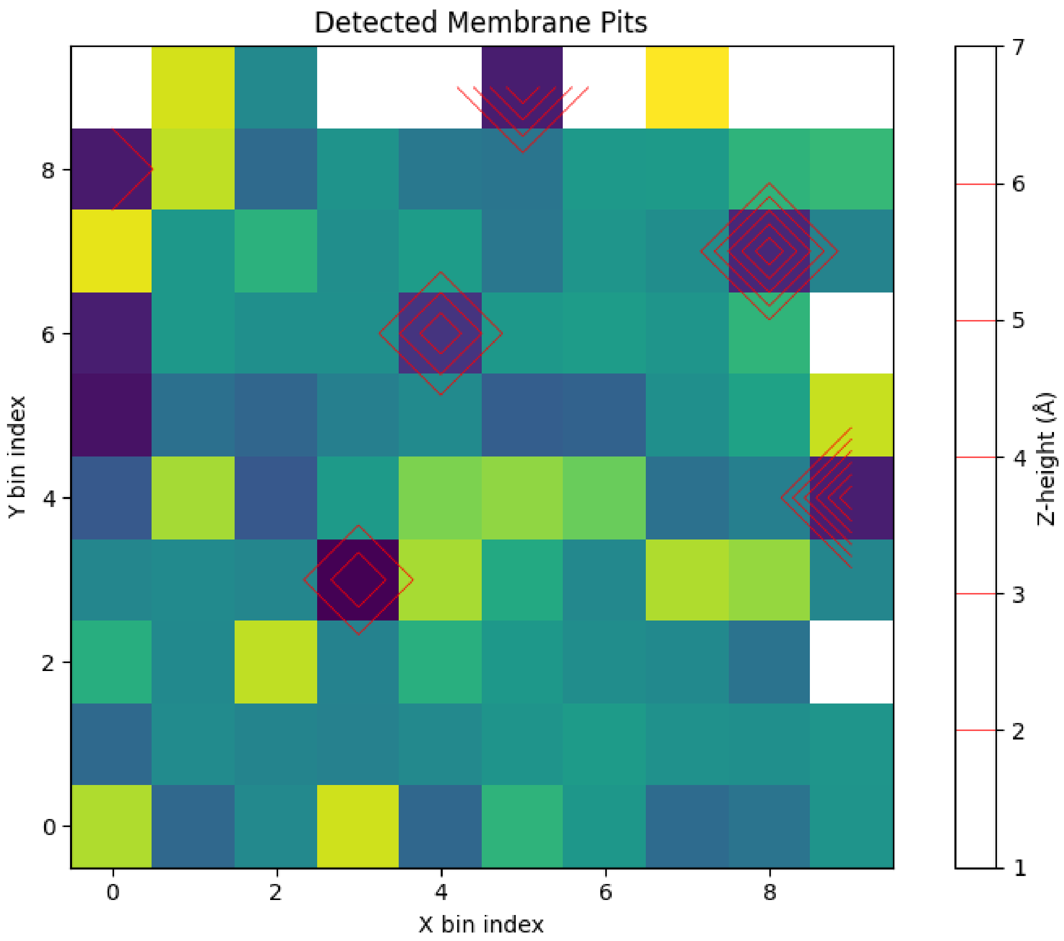


**Supplementary Figure S7. Topographical map of membrane surface with detected pit regions.** A 2D heatmap of the membrane Z-height is shown, representing local membrane elevations and depressions across a protein-embedded lipid bilayer. Z-height values (in Å) are colored using the *viridis* colormap, with darker shades indicating lower membrane regions. Red contours mark the boundaries of connected pit regions, defined as contiguous bins where Z-height values fall ≥1.5 standard deviations below the mean surface height. Each pit is labeled and corresponds to localized deformations in the membrane topology. These regions may reflect protein-lipid interactions, curvature induction, or hydrophobic mismatch. The colorbar denotes Z-height in Ångströms.

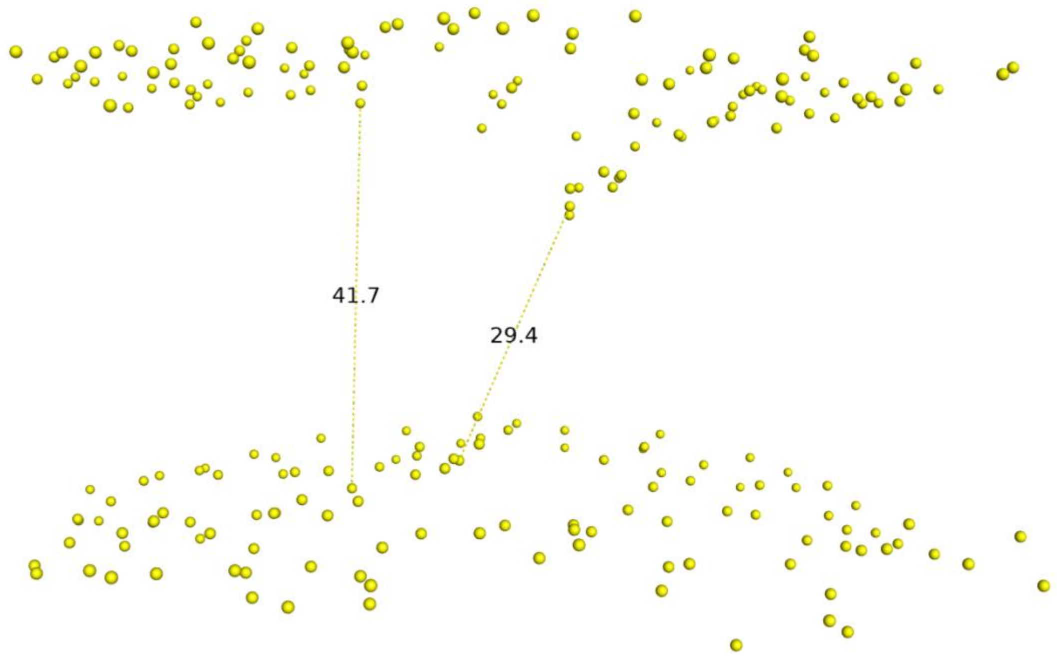


**Supplementary Figure S8. Lipid head phosphates of the membrane-protein system after 250ns full-atom MD simulation.**
